# Methylmalonic acid compromises mitochondrial respiration and reduces the expression of markers of differentiation in SH-SY5Y human neuroblastoma cells

**DOI:** 10.1101/2020.08.24.265157

**Authors:** Renata Torres da Costa, Marcella Bacelar dos Santos, Izabel Cristina Santos Silva, Raquel Pascott Almeida, Marcela Simões Teruel, Daniel Carneiro Carrettiero, César A. J. Ribeiro

**Author notes:** Corresponding Author Cesar. A. J. Ribeiro, PhD, Universidade Federal do ABC – UFABC, Centro de Ciências Naturais e Humanas – CCNH, Rua Arcturus, n° 03, Bloco Delta, sala 231, Jardim Antares, São Bernardo do Campo, SP, CEP 09606-070, Brazil.

## Abstract

Methylmalonic acidemia is a rare metabolic disorder characterized by the accumulation of methylmalonic acid (MMA) and alternatives metabolites which is caused by the deficient activity of L-methylmalonyl-CoA mutase or its cofactor 5-deoxyadenosylcobalamin (AdoCbl). The brain is one of the affected tissues by the accumulation of this metabolite in patients. The neurologic symptoms commonly appear in newborns and are clinically characterized by seizures, mental retardation, psychomotor abnormalities, and coma. The molecular mechanisms of neuropathogenesis in methylmalonic acidemia are still poorly understood, specifically regarding the impairments in neuronal development and maturation. In this study, we firstly investigated the neurotoxicity of MMA in both undifferentiated and 7-day RA-differentiated phenotypes of SH-SY5Y human neuroblastoma cells and found alterations in energetic homeostasis after the exposition to MMA. We observed an increase in glucose consumption and reduced respiratory parameters of both undifferentiated and differentiated SH-SY5Y cells after 48 hours of exposition to MMA. RA-differentiated cells slightly indicated to be more prone to perturbations in respiratory parameters by MMA than undifferentiated cells. In order to understand whether the presence of MMA during neuronal maturation could compromise this process in neuronal cells, we performed high-resolution respirometry to evaluate the mitochondria function and qPCR assay to evaluate mRNA levels of mature neuronal-specific genes in early-stage (day 3), and late-stage (day 7) of differentiation in cells co-treated with MMA 1mM during RA mediated differentiation. Our results showed that MMA compromises the respiratory parameters of routine, ATP-linked, and maximal respiration only at the late stage of differentiation as well as downregulates the transcriptional gene profile of mature neuronal markers ENO2 and SYP. Altogether, our finds point to important alterations observed during neuronal maturation and energetic stress vulnerability that can play a role in the neurological clinical symptoms at the newborn period and reveal important molecular mechanisms that could help the screening of targets to new approaches in the therapies of this disease.

## 1. INTRODUCTION

Inborn errors of metabolism are rare genetic disorders that comprise several groups of disturbs, classified according to the affected metabolism pathway or accumulating product. Among them, methylmalonic acidemia, characterized by the accumulation of methylmalonic acid (MMA) and alternatives metabolites, is caused by the deficient activity of L-methylmalonyl-CoA mutase or its cofactor 5-deoxyadenosylcobalamin (AdoCbl) [1,2]. Despite the diverse phenotypes and etiologies of mutations described for methylmalonic patients, the first clinical symptoms are similar and commonly appear in newborns. At disease onset, the common symptoms are lethargy, vomiting, and refusing to feed; in severe cases, patients can evolve to neurologic manifestations as seizures, mental retardation, psychomotor abnormalities, and coma [1].

MMA is produced during the propionate metabolism which takes place inside the mitochondrial matrix. During a patient metabolic crisis, this organelle is more sensitive to alterations in its functionality due to the high concentration of MMA inside the mitochondria than in the cytosolic compartment of the cells [3]. The accumulation of methylmalonic acid is proposed to cause complications in specific organs among them the brain [4].

Neurologic dysfunctions in methylmalonic acidemia are important clinical manifestations in patients. Previous studies suggested that primary or secondary mitochondrial dysfunction is potentially elicited by MMA, proposing that these dysfunctions are related to the neurological alterations noticed in patients [4,5]. Moreover, it has been argued that other metabolites derived from the accumulation of MMA may also be the factors responsible for dysfunction in mitochondrial activity [7–9].

The neuropathogenesis noticed in methylmalonic acidemia patients are still poorly understood as well as the levels of MMA inside the brain tissue are not well characterized. Cerebrospinal fluid values are used as estimates for neuronal levels of MMA [10,11], however, it has been discussed in the literature that the presence of the blood-brain barrier may contribute to trapping these compounds inside of the brain, reaching high concentrations of MMA and inducing neurological complications in patients [7]. The nervous system presents a high energy metabolism demand compared to other tissues, among the cells presented in this tissue, neurons cells display the highest energy requirements that can be altered during the neuronal development [12]. Regarding the pivotal role of mitochondria in neuronal activity and development, this study aimed to investigate the effects of MMA in bioenergetics and neuronal development in human neuroblastoma cells line which presents neuronal features. We evaluated routine, ATP-linked, leak, maximal respiration, spare respiratory capacity, and non-mitochondrial respiration.

## 2. MATERIAL AND METHODS

### 2.1. Reagents

Fetal bovine serum (FBS), penicillin/streptomycin, and trypsin were purchased from Gibco (Carlsbad, CA, USA). SY-SY5Y human neuroblastoma cells were obtained from Banco de Células do Rio de Janeiro. Methylmalonic acid, Dulbecco’s modified Eagle’s medium (DMEM, containing 5.5 mM glucose), methyl-thiazolyl diphenyl tetrazolium bromide (MTT), as well as all other chemicals of analytical grade were obtained from Sigma (St Louis, MO, USA). Methylmalonic acid and other reagents were acquired from Sigma Aldrich, all-trans-retinoic acid from Merck Millipore. Probes for qPCR were purchased from in IDT DNA (Integrated DNA Technologies Inc, USA)

#### a. Cell culture

Human neuroblastoma cells (SH-SY5Y) were cultivated in low glucose Dulbecco’s Minimum Essential Medium (DMEM), supplemented with 10% v/v fetal bovine serum and 100 U/mL of penicillin and 100 μg/mL of streptomycin. The SH-SY5Y cells were grown to confluence at 37°C under 5% CO_2_ atmosphere and maintained between passages 10 to 16. The cells were seeded for experiments at 1,5 × 10^5^ cells/cm^2^. Experiments were done with undifferentiated and differentiated cells. All reagents used were dissolved in their respective vehicles (growth medium, DMSO, ethanol, or phosphate buffer saline).

#### b. SH-SY5Y Exposition to Methylmalonic Acid

Methylmalonic acid (MMA) was dissolved in PBS, adjusted pH to 7.4, and sterilized by filtration in 22 μm membrane, in the day of the experiment. SH-SY5Y cells were exposed to MMA (final concentrations of 0.1, 1.0, 5.0, and 10.0 mM) in DMEM medium supplemented with 5% FBS for 24 or 48 hours. Cells in the control group were maintained in DMEM with 5% FBS, supplemented with the same volume of PBS used in MMA-exposed cells.

#### c. Differentiation of SH-SY5Y cells

Experiments were performed with undifferentiated and retinoic acid (RA)-differentiated SH-SY5Y cells. For differentiation, SH-SY5Y cells were seeded at 0,6 × 10^5^ cells/cm^2^, and 24 hours after plating cells were treated with 10 μM of all-trans-retinoic acid (RA) for 7 days in DMEM medium containing 1% FBS. The medium was renewed every 72-hour. This differentiation protocol is known to induce a mature dopaminergic-like neuronal phenotype [13]. Additionally, to evaluate the toxic effects of MMA during neuronal maturation, SH-SY5Y cells under 10 μM of RA-differentiation treatment were co-incubated with 1 mM of MMA during 3 days (early stage of differentiation) and 7 days (final stage of differentiation).

#### d. Cytotoxicity and glucose consumption assays

Cytotoxicity caused by MMA exposition was evaluated by MTT (3-(4,5-dimethylthiazol-2-yl)-2,5-diphenyltetrazolium bromide) reduction assay. For this assay SH-SY5Y cells were seeded at properly amounts for undifferentiated and RA-differentiated cells in 96-well plate. After the exposition to MMA for 24 or 48-hour periods, the cells were incubated with 0.5mg/mL of MTT solution for 2 hours at 37°C. Formazan crystals were solubilized by adding isopropanol. Absorbance was measured at 570nm in a plate reader EPOCH Biotek. Glucose consumption assay was performed using a commercial kit based on glucose oxidase (Labtest, Ref.133 – Minas Gerais, Brasil). SH-SY5Y cells supernatants were collected and prepared according to the manufacturer’s instructions and incubated by 10 minutes at 37°C. Absorbance was measured at 550nm in a plate reader EPOCH Biotek. Basal levels of glucose in the DMEM medium before exposition to cells were used to calculate the amount of glucose consumption during the period of exposition to MMA.

#### e. Characterization of mitochondrial respiration with oxygen consumption measurements

Oxygen consumption in undifferentiated and RA-differentiated SH-SY5Y cells was measured by high-resolution respirometry using Oroboros O2K Oxygraph (OROBOROS Instruments, Innsbruck, Austria) in 2 mL chamber at a stirrer speed of 750 rpm. After exposition to MMA, cells were detached and counted. Oxygen consumption in both undifferentiated and RA-differentiated SH-SY5Y cells was determined with 3 × 10^6^ cells per chamber in CO_2_ equilibrated DMEM medium with 5% FBS. Manual titration of inhibitors and uncoupler was performed using Hamilton syringes (Hamilton Company, Reno, NV, USA). The following inhibitors and uncoupler (final concentrations) were added to access the different respiratory states: 0.5 μM of oligomycin to inhibition of ATP-synthase, 2.5 μM of CCCP as a mitochondrial uncoupler, 0.5 μM of rotenone to inhibit mitochondrial complex I and 0.5 μM of antimycin A, an inhibitor of mitochondrial complex III. To investigate the mitochondrial respiration, the following respiratory parameters were calculated: (I) Routine respiration (basal oxygen consumption employed to supply cellular regular metabolism), (II) proton leak respiration (oxygen consumption related to proton flux across the mitochondrial inner membrane), (III) ATP-linked respiration (amount of oxygen consumed by cells to produce ATP, determined by the rate of respiration that can be inhibited by oligomycin), (IV) Maximal respiration (oxygen consumption after mitochondrial uncoupling, related to maximal electron flux in the electron transfer chain), (V) Spare respiratory capacity (the difference between maximal and routine respiration) and (VI) non-mitochondrial respiration (ROX, related to oxygen consumption after inhibition of complexes I and III of the electron transfer chain). Data recording was performed using the DataLab software 7.04 (OROBOROS Instruments). Oxygen flux in all the respiratory parameters was corrected for non-mitochondrial respiration (ROX) and expressed as pmol O_2_ / s / 10^6^ cells.

#### f. Analyses of mRNA expression by qPCR assay

The effect of MMA exposition during neuronal development was evaluated by molecular markers of mature neuronal phenotype in SH-SY5Y cells. Total RNA was isolated from undifferentiated cells, RA-differentiated cells, and RA-differentiated cells + 1.0 mM of MMA groups by using Tri Reagent®. RNA integrity and quality were confirmed by BioDrop μLITE® Spectrophotometer (Cambridge, United Kingdom). cDNA was transcribed using the High Capacity cDNA Reverse Transcription (Applied Biosystems). Quantitative PCR (qPCR) reactions using TaqMan chemistry were performed with probes for ENO2 and SYP (Hs.PT.58.578449 and Hs.PT.58.272077712, respectively, purchased from Integrated DNA Technologies Inc.) on a StepOne Real-Time PCR System and Applied Biosystems StepOne software 2.0 (Foster City, USA). The comparative cycle threshold (Ct) method was used to determine the relative quantity of mRNA and amplification efficiency and sample to sample variation were normalized with Hypoxanthine-guanine phosphoribosyltransferase (HPRT) as an internal control gene.

#### g. Statistical analyses

Data are expressed as mean ± SEM (n=4-6 as described in figure legends). Comparisons between two groups were performed with unpaired *Student t*-tests. Comparisons between multiple groups were performed with One-way or Two-way ANOVA with Tukey’s post hoc analyses for comparisons between control and treatment groups. A *P* value of less than 0.05 was considered significant. All the statistical analyses were performed with Prisma Software (version 6.07, GraphPad Software Inc, San Diego, CA).

## 3. RESULTS

### 3.1. Exposition to retinoic acid promotes alterations in oxygen consumption and gene expression profile between undifferentiated and RA-differentiated SH-SY5Y cells

The effects of RA on the expression of molecular markers of mature neuronal phenotype, namely synaptophysin (SYP) and neuron-specific enolase (ENO2), were evaluated in undifferentiated (day 0) and RA-differentiated SH-SY5Y cells (at days 3 and 7) by qPCR assay. As shown in Figure 1A, RA-induced differentiation of SH-SY5Y cells was confirmed by a significant increase in the expression of ENO2 gene after 3 and 7 days of RA treatment when compared to undifferentiated SH-SY5Y cells (*P* < 0.05). The expression of SYP was significantly elevated at day 3 and day 7 of RA treatment compared to undifferentiated SH-SY5Y cells gene (*P*<0.001, Figure 1B), reaching the highest levels at day 7. Additionally, morphologic alterations after exposition to retinoic acid were observed after 3 days of starting the RA differentiation treatment in SH-SY5Y cells and remained without differences until day 7 (Figure 1C). These data validate the progress and differentiation of SH-SY5Y cells via neuronal maturity markers, after exposition to RA treatment for 7 days.

**Figure 1.**
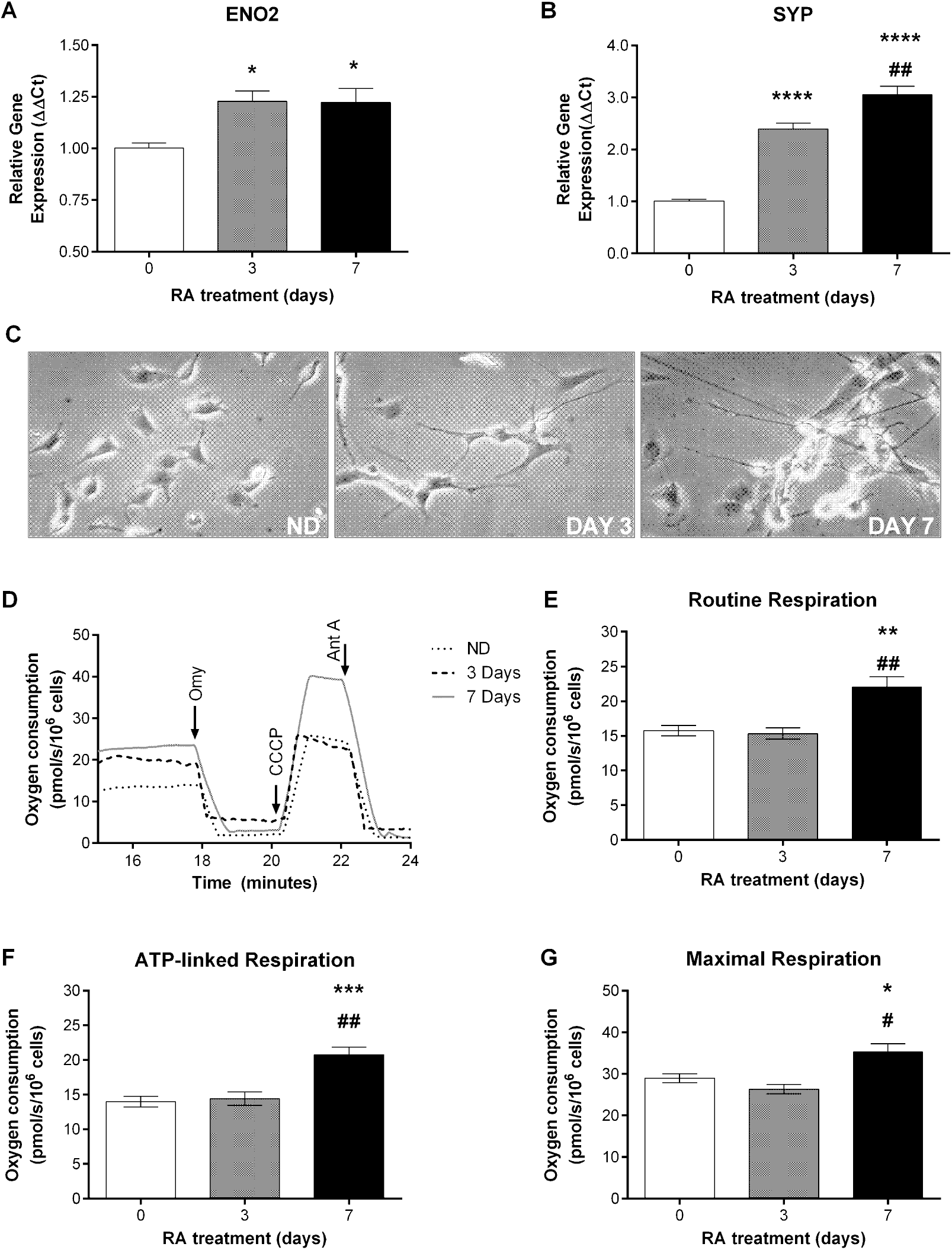
Alterations in the expression of neuronal mature markers neuron-specific enolase (ENO2) and synaptophysin (SYP) and increase routine, ATP-linked and, maximal respiration after prolonged exposition to retinoic acid treatment. Quantification of mRNA expression for (A) ENO2 and (B) SYP was increased during RA treatment in SH-SY5Y cells, confirming a mature neuronal phenotype. Oxygen consumption in undifferentiated (day 0) and RA-differentiated SH-SY5Y cells (days 3 and 7) was measured by high-resolution respirometer Oroboros. (C) Morphologic changes in SH-SY5Y during RA-differentiation. Cells were visualized with a 20x objective. To determine different parameters of mitochondrial function, (D) oligomycin, an ATP synthase inhibitor (0,5μM), uncoupler CCCP (2,5μM), rotenone, a complex I inhibitor (0,5μM) and antimycin A, complex III inhibitor (0,5μM) were injected in the chamber with cells. Routine respiration (E) was determined before injected oligomycin, ATP-linked respiration (F) was calculated as the fraction of oxygen consumption inhibited by oligomycin. Maximal respiration (G) was measured after the injection of CCCP. Statistical analysis performed with one-way ANOVA and multiple comparisons with Tukey’s post-hoc test. n=5 for each experiment. Data are expressed as mean ± SEM. * p<0.05, ** p<0,01, *** p<0.001 compared between day 0 and day 7 samples. # p<0.05, ## p<0.01 compared between day 3 and day 7.

In order to analyze possible changes in cellular bioenergetics after RA treatment, we access the cell respiration in undifferentiated and RA differentiated SH-SY5Y cells using high-resolution respirometry. Polarographic measures of oxygen consumption were performed in SH-SY5Y without unexposed to RA (day 0) as well as in cells exposed to RA for 3 (early stage of differentiation) and 7 days (late stage of differentiation). Figure 1C shows representative experiments of each experimental group and the additions inhibitors and uncoupler in the course of the assay. Regarding routine respiration, it was noticed a significant increase only in 7-day RA-differentiated SH-SY5Y cells (p<0.01, Figure 1D) when compared to undifferentiated and 3 days RA-differentiated groups. It was also observed a significant increase in oxygen consumption linked to ATP production (ATP-linked respiration) and the maximal respiration only in 7-day RA-treatment SH-SY5Y cells (Figure 1, panels E and F), suggesting that changes in the cellular bioenergetics occur only in the late stages of differentiation with retinoic acid.

### 3.2. Metabolic viability decreased in undifferentiated and 7-day RA-differentiated SH-SY5Y cells after exposition to MMA

In order to elucidate the possible effects of MMA in the metabolism of a non-mature and mature neuron, we performed assays to investigate metabolic cytotoxicity and glucose consumption in undifferentiated and 7-day RA-differentiated SH-SY5Y cells after exposition to distinct concentrations of MMA. We observed a significant increase in glucose consumption 24 and 48-hour after cell exposition to MMA, and this effect was much more pronounced in undifferentiated than RA-differentiated cells (*P* <0.05, Figure 2, panels A and B). Interestingly, MMA reduced metabolic viability in both undifferentiated and RA-differentiated cells after 48-hour of exposition in a dose-dependent manner (*P*<0.05, Figure 2, panels C and D). Undifferentiated and RA-differentiated SH-SY5Y cells exposed to MMA showed differences in glucose consumption and metabolic viability. Together, these results suggest an impairment of metabolic viability of SH-SY5Y cells after exposition to MMA and it’s corroborated by the alterations observed in glucose levels consumption.

**Figure 2.**
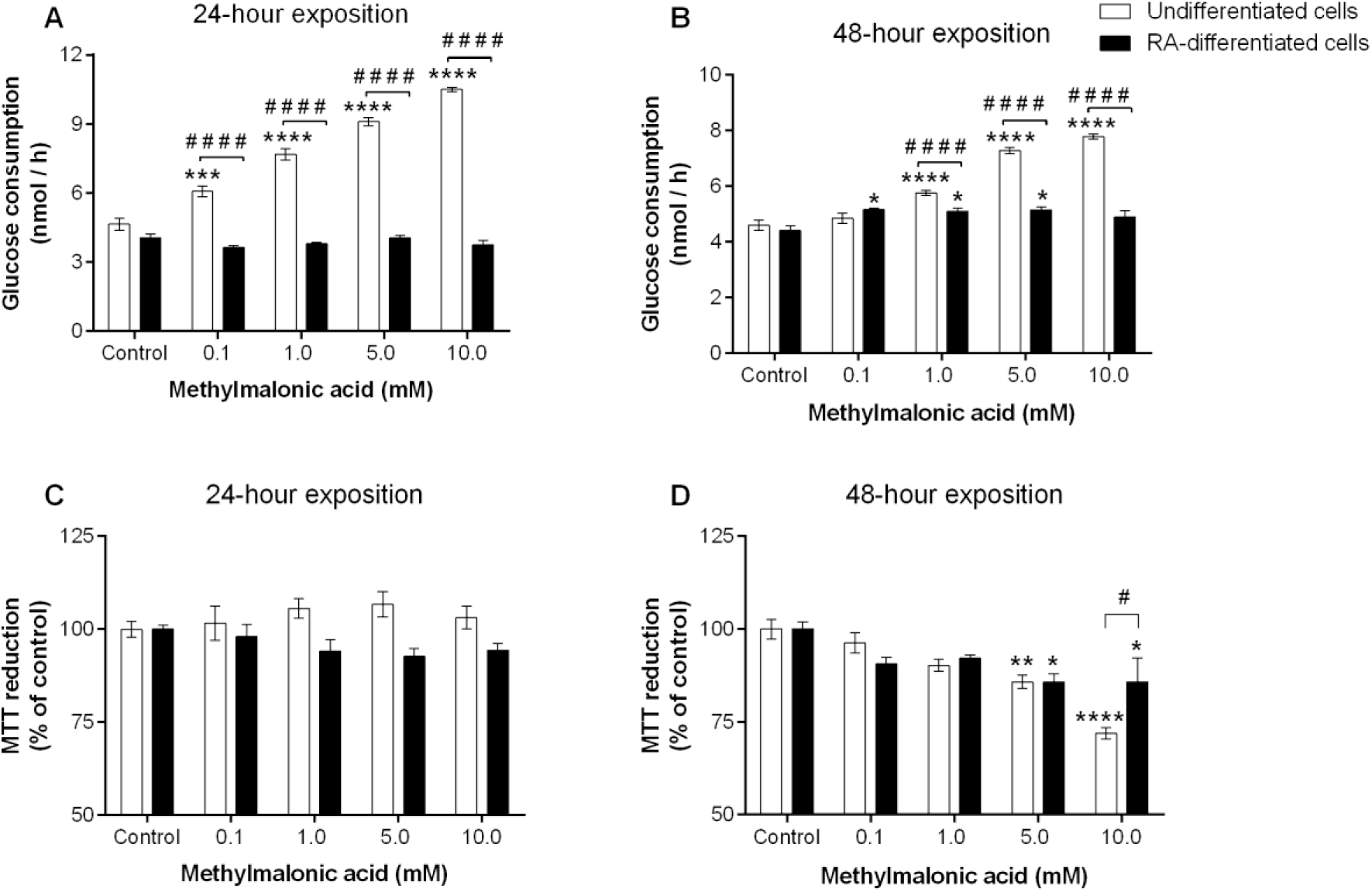
Undifferentiated SH-SY5Y cells are more prone to increase glucose consumption and reduce metabolism viability after MMA exposition. Glucose consumption of undifferentiated and 7-day RA-differentiated SH-SY5Y cells after (A) 24-hour and (B) 48-hour exposition to MMA. MTT reduction assay of undifferentiated and 7-day RA-differentiated SH-SY5Y cells after exposition to MMA for (C) 24 and (D) 48 hours. Data = mean ± SEM. * p<0.05, ** p<0.01, ***p<0.001, ****p<0.0001. n=4 for each experiment. Statistical analyses were carried out with two-way ANOVA, Tukey’s post-hoc test compared to the respective control group. # p<0.05, #### p<0.0001 compared to the related RA-differentiated group.

### 3.3. Methylmalonic acid compromises energy metabolism reducing oxygen consumption in undifferentiated and 7-day RA-differentiated SH-SY5Y cells

As our results suggested a possible impairment in cellular bioenergetic homeostasis caused by MMA, we next investigated whether MMA alters respiratory states in both undifferentiated and 7-day RA-differentiated SH-SY5Y cells. We investigated the mitochondrial respiratory states of these cells after 48-hour exposition to MMA (1.0, 5.0, and 10.0 mM).

As shown in Figure 3A, routine respiration was significantly reduced in undifferentiated and RA-differentiated SH-SY5Y cells in response to MMA exposition in concentrations of 5.0 and 10.0 mM. Next, we accessed the oxygen consumption linked to ATP (ATP-linked respiration, Figure 3B) which was significantly reduced in a dose-response manner at RA-differentiated cells. However, effects on this parameter in undifferentiated cells were only observed in the highest concentration of MMA tested (10mM). Regarding the maximal respiration and spare respiratory capacity of both cell conditions, it was noticed a significant reduction of about 30-40% in these parameters to respective control groups (Figure 3, panels C and D). No significant differences were observed in comparisons between the groups of undifferentiated and RA-differentiated cells exposed to MMA at the same concentrations.

**Figure 3.**
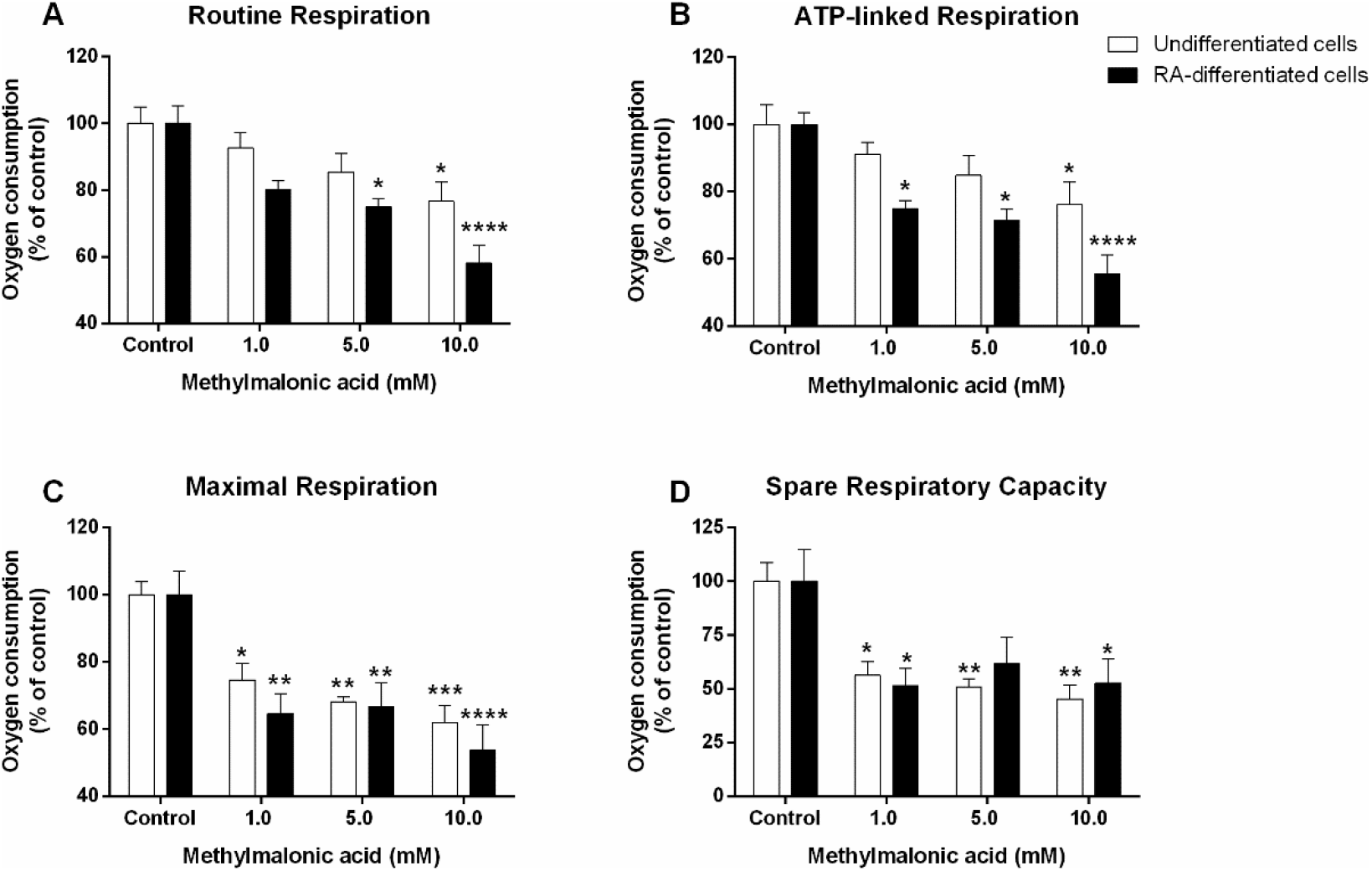
Effects of methylmalonic acid in the respiratory parameters of undifferentiated (day 0) and RA-differentiated (day 7) SH-SY5Y cells after 48 hours exposition. Cells were exposed to three concentrations of MMA (1.0, 5.0, and 10.0 mM) for 48 hours and measured the oxygen consumption in both conditions: undifferentiated and 7-day RA-differentiated SH-SY5Y cells and, the following respiratory parameters were determined: (A) routine respiration, (B) ATP-linked respiration, (C) maximal respiration and, (D) spare respiratory capacity. The oxygen consumption was expressed as a percentage of the respective controls group. Data= mean ± SEM from 6 independent experiments. Statistical analysis performed with two-way ANOVA, multiple comparisons by Tukey’s post-hoc test. * p<0.05, ** p<0.01, *** p<0.001, ****p<0.0001 compared to the respective control groups.

These results imply that MMA altered the function and control of the electron transport chain and ATP production in both undifferentiated and RA-differentiated SH-SY5Y cells. Regardless of no significant differences were noticed at respiratory states between the non-mature and mature neuronal phenotypes of SH-SY5Y cells exposed to MMA, RA-differentiated SH-SY5Y seems to be slightly more prone to significant mitochondrial dysfunctions after exposition to MMA, suggesting a higher susceptibility of mature neurons to alterations in mitochondrial function.

### 3.4. Exposition to MMA during differentiation compromises mitochondrial function and mRNA expression profile at the late stage of RA-differentiation process in SH-SY5Y cells

We also evaluated the effects of MMA on the neuronal maturation process. In these experiments, cells were co-incubated with MMA (1 mM) and RA (10 μM), and the respiratory parameters, as well as the expression of SYP and ENO2 genes, were analyzed at early (3 days) and late stage of differentiation (7 days).

As shown in Figure 1 (panels D, E and, F), 7-day exposition to RA significantly increased the respiratory parameters of cells compared to undifferentiated SH-SY5Y cells. Interestingly, the respiratory parameters of 7-day RA-differentiated cells were compromised by co-exposure to 1 mM MMA Figure 4 A). Oxygen consumption in routine, ATP-linked, and maximal respiration was significantly reduced (about 20-30%) in RA plus MMA-exposed cells when compared to RA alone cells (control group). No significant differences were observed at the co-exposure with MMA in 3-day RA-differentiated SH-SY5Y cells and some of the respiratory parameters remained similar to values of undifferentiated cells. Altogether, these results suggest that SH-SY5Y cells in the late stage of the differentiation are more vulnerable to dysfunction on mitochondrial respiratory parameters than the SH-SY5Y cells in the early stage of this process.

**Figure 4.**
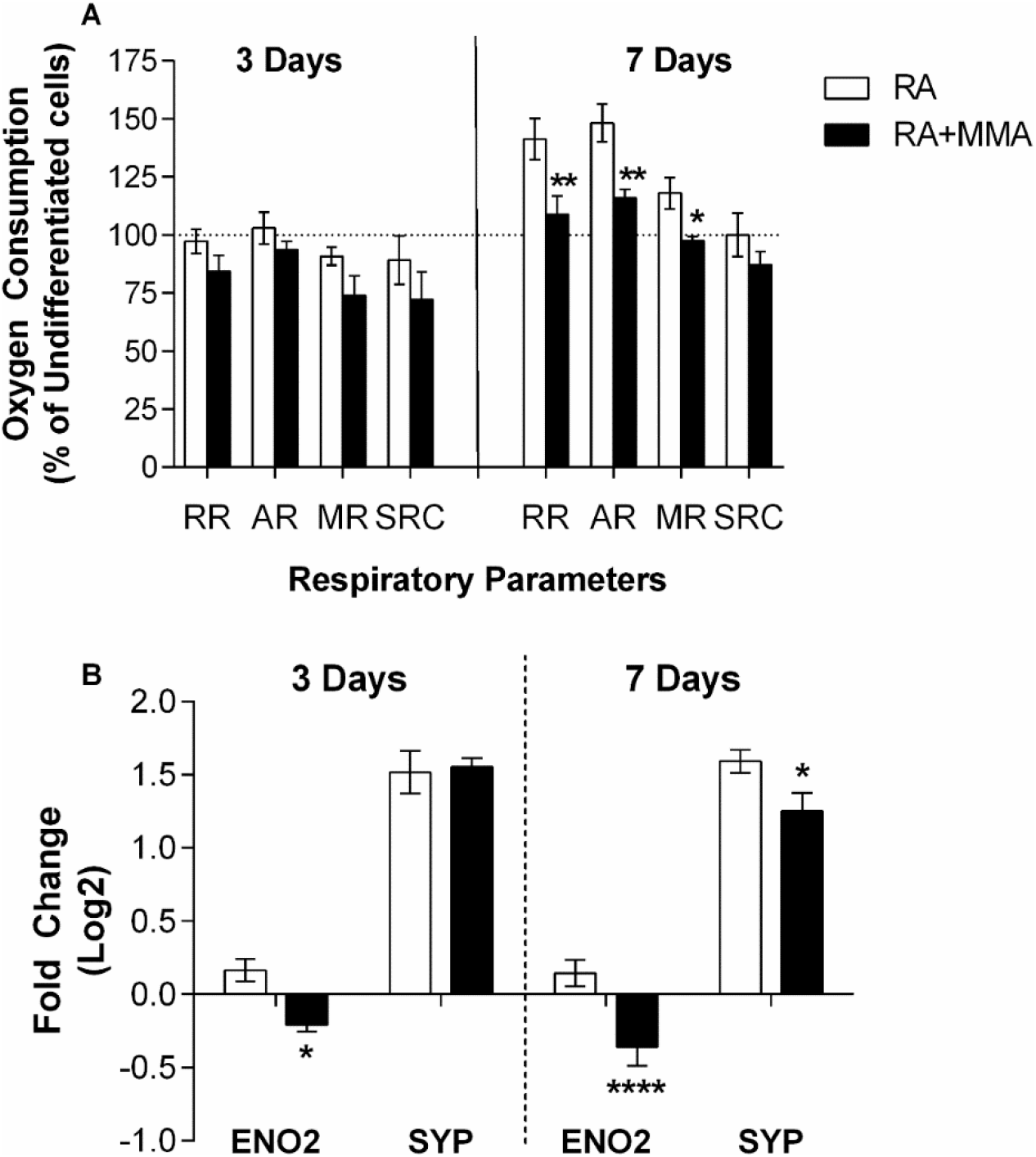
Comparisons of MMA effects in respiratory parameters and mRNA expression levels of neuronal mature markers in 3-day RA-differentiated and 7-day RA-differentiated SH-SY5Y cells. (A) Oxygen consumption levels were measured by high-resolution respirometry assay in both stages of the RA differentiation process: early (day 3) and late (day 7) stages at SH-SY5Y cells exposed to MMA 1mM (RA+MMA) and respiratory parameters were determined. RR: routine respiration, AR: ATP-linked respiration, MR: maximal respiration and, SRC: spare respiratory capacity. The oxygen consumption is expressed as the percentage of undifferentiated SH-SY5Y cells group. The dashed line represents the percentage of undifferentiated SH-SY5Y cells. (B) A comparative profile of mRNA expression levels of 3-day RA differentiated and 7-day RA-differentiated SH-SY5Y cells co-exposed to MMA 1mM during the maturation process. The levels of expression are shown on a Log2 scale as fold change, compared to the respective levels of mRNA in undifferentiated cells. Bars are mean ± SEM from 6 independent experiments. Statistical analysis was carried out with two-way ANOVA, multiple comparisons by Tukey’s post-hoc test. * p<0.05, ** p<0.01, ****p<0.0001 compared to the respective control groups.

We also investigated whether the presence of MMA during RA-induced differentiation could compromise the expression of mature neuronal markers at the different stages of neuronal maturation. Co-incubation of cells with MMA and RA for 3 days significantly reduced mRNA expression for ENO2 gene (Figure 4 B). The presence of MMA during the 7 days (late-stage) of RA-differentiation in SH-SY5Y cells significantly reduced mRNA levels in both ENO2 and SYP genes (Figure 4B), indicating that MMA affected the profile of these neuronal markers.

## 4. DISCUSSION

The central nervous system (CNS) is one of the affected tissues by the accumulation of MMA during the metabolic crisis in methylmalonic acidemia patients [14]. The molecular mechanisms of neurologic impairments in methylmalonic acidemia are still poorly understood but the development of new models and approaches for the study of neurodegeneration are important tools to elucidate these events. In this regard, the human neuroblastoma SH-SY5Y cell line is widely used as an *in vitro* model for the investigation of several neurodegenerative disorders, such as Alzheimer’s [15], Parkinson’s [16], and Huntington’s disease [17], demonstrating to be a reliable study model and highlighting the potential applications of this cell line in organic acidemias investigations. SH-SY5Y cells can present two distinct profiles: undifferentiated cells, characterized by the lack of mature neuronal properties, and, when exposed to RA combined with lowering FBS, SH-SY5Y cells give rise to the neuronal-like phenotype, presenting morphological and biochemical characteristics of a mature neuron [16,18]. Regarding the two profiles of SH-SY5Y cells, in this study, we explored these properties as a novel approach to investigate the neurotoxicity of MMA in distinct stages of neuronal development. First, we evaluated the response of SH-SY5Y to RA exposition during two stages of the differentiation process (days 3 and 7). Both neuron-specific markers ENO2 and SYP were increased after exposition to RA on day 3 and reaching the high levels on day 7, which are in agreement with previous studies reported in the literature [18], and validate the differentiation process since the expression of these neuron-specific genes is related to neuronal maturation.

Moreover, considering that cellular bioenergetics dynamics of neurons can be reprogrammed during neuronal development [19], it’s important to clarify possible changes that RA differentiation can promote in SH-SY5Y cells. Supporting this notion, mitochondrial function was assessed in both conditions: undifferentiated and RA-differentiated cells. Our results indicated that RA-differentiation significantly increases oxygen consumption in routine, ATP-linked, and maximal respiration only in the late stage (day 7) of differentiation in SH-SY5Y cells. The lack of alterations in these respiratory parameters at the early stage (day 3) of differentiation suggests that changes in the cellular bioenergetics may occur in the advanced phases of neuronal development. Indeed, it’s known that after neuronal maturation, cells present a higher energy demand to support the new functionalities acquired from this process [20]. In support of that, oxygen consumption was found to increase specifically in the respiratory parameters associated with ATP production and the maximal capacity of the electron transfer chain. It was also found at the late stage of differentiation an increase in the expression of the SYP gene, which is related to the synaptic process, reflecting a functional maturity in neurons [21]. In previous reports, similar results were found regarding the increase of mitochondrial respiratory parameters, as maximal respiration and bioenergetics reserve capacity [22].

The accumulation of MMA is known to cause dysfunction in neuronal development which has been related to the clinical symptoms in patients [6,23]. However, there is a lack of investigations about the molecular mechanisms involved in the alterations in neuronal differentiation of immature and mature neurons. In that sense, we examined the toxic effects of MMA on undifferentiated and differentiated SH-SY5Y cells. First, we evaluated the cellular viability of both profiles of SH-SY5Y cells after exposition to MMA. As shown in Figures 2A and 2B, undifferentiated SH-SY5Y cells significantly increased glucose consumption in both periods of exposition to MMA in a dose-dependent manner. Although 7-day RA differentiated cells also showed an increase in glucose consumption compared to the respective control group, the effects of MMA in this parameter showed to be not so remarkable as the results observed in undifferentiated cells. More likely, these findings may be sharper in undifferentiated cells mainly because of the contrast in the cellular bioenergetic requirements that this SH-SY5Y phenotype presents compared to the RA-differentiated, as reported in previous studies [24].

Altogether, these findings suggest a possible mitochondrial impairment in both cell conditions since the mitochondrial based reduction of MTT was reduced after exposition to MMA. In this particular, to compensate for the dysfunction in mitochondria respiration, more glucose is utilized to supply cell energy demand. Neurons can suffer more than other cells from CNS since their energy production is considerably supported by oxidative phosphorylation raising serious consequences for brain energy metabolism [25,26]. Moreover, MMA increased glucose consumption, suggesting that undifferentiated and RA-differentiated SH-SY5Y cells may be employing more glycolysis than oxidative phosphorylation to supply their ATP demand in response to perturbations in mitochondria function caused by the high concentrations of MMA. Previous reports showed hypoglycemia and lactic acidosis as a particular biochemical remark in this organic acidemia, suggesting dysfunction in energy metabolism [27,28].

These different effects observed on cell viability and glucose consumption raised the question of whether immature or mature neuronal cells would be more vulnerable to mitochondrial dysfunction caused by MMA. In this regard, we observed that RA-differentiated cells seem to be slightly more susceptible to mitochondrial alterations at lower concentrations of MMA than undifferentiated cells. Routine and ATP-linked respiration was reduced in the presence of MMA, suggesting a direct inhibitory effect on oxidative metabolism or a perturbation in substrates supply for the mitochondria. Interestingly, maximal respiration was significantly reduced in both cell conditions, reinforcing the role of MMA in mitochondrial dysfunction. Moreover, we also reported a significant decrease in reserve respiratory capacity in both undifferentiated and differentiated cells exposed to all concentrations of MMA. Reserve respiratory capacity would be employed for distinct purposes depending on the cell phenotype. In non-proliferative cells, as mature neurons, it can be used to supply energy requirements to survive stress. In contrast to that, proliferative cells will use most of the reserve capacity to perform the mitotic process [29]. We also reasoned that RA-differentiated cells may be more prone to MMA toxicity since the present results indicated that cells at the late stage of differentiation (day 7) had an increase in ATP linked respiration suggesting a preference for oxidative phosphorylation pathway which could reflect a vulnerability to energetic stress related to mitochondrial dysfunctions.

Although we cannot correlate observed alterations on cell oxygen consumption to an inhibitory effect of MMA since we did not evaluate the mitochondrial complexes activity in this present study, it was demonstrated in different experimental models that MMA disrupts mitochondrial energy metabolism [5,30], by inhibiting electron transfer chain complexes I [5,31], II [32], and compromises the transport of substrates by di-carboxylic transporters to the mitochondria [30,32]. On the other hand, some studies pointed out that mitochondrial dysfunction is not mainly caused by MMA, but impairment of mitochondrial function is likely to be related to the secondary metabolites that can be accumulated in methylmalonic acidemia [8].

In the context of development and maturation of the nervous system, the first few years of life are considered the most important period [33], and the first metabolic crisis of methylmalonic acidemia patients usually occurs during this stage of life [1]. In order to understand the possible neurotoxicity of MMA during the neuronal maturation process, we evaluated if MMA compromises respiration during cellular differentiation by co-exposing of cells to RA and MMA. Late differentiated cells (day 7) showed a significant decrease in routine, ATP-linked, and maximal respiration. The lack of effects in the early stage of differentiation reflects the distinct energetic dynamics between these two stages of neuronal maturation. As shown in our previous results, the early stage of differentiation by RA exposition provides an indicator that cellular bioenergetic is similar to the undifferentiated cells which depend on the glycolytic pathway. More likely, the bioenergetics shift observed in the late stage of differentiation, reported in previous studies [34], corroborates to increase the sensibility of these cells to mitochondria perturbations.

The development of the mammalian brain is regulated by several mechanisms including transcriptional gene regulation and expression, in order to accomplish the morphological and functional alterations for the maturation of neurons [35]. Since during these changes mitochondria play an important role in providing the necessary ATP level to neuron cells [12,36], we also evaluated whether mitochondrial dysfunctions caused by MMA during the differentiation process would compromise the gene expression profile of mature neuronal markers. Quantification of mRNA levels of mature neuron markers by qPCR confirmed the changes in the expression profile on these cells caused by MMA.

Our finds indicated that in the early stage of differentiation, MMA decreased mRNA levels of ENO2, in contrast to late-stage (day 7) where both SYP and ENO2 were reduced. In this regard, prior studies have shown an essential role of mitochondria during brain development in embryonic, postnatal, and even in the adult period indicating that perturbations could lead to neurological manifestations related to cognitive abnormalities [37,38]. The reduction observed in the expression of these markers of mature neurons in our experimental model indicates that the presence of MMA during the differentiation process caused a delay in the maturation most pronounced in the late stage which could cause perturbations in the functionality of neuronal cells.

Prior studies in the literature employing the differentiation of immature cells and mature neurons have revealed that a decrease in the expression levels of molecular markers of this process is related to cognition abnormalities [39]. The use of SH-SY5Y cells in different stages of the RA-differentiated process consists of a promising model system to study the role of mitochondria dysfunction on the onset of neurological symptoms in methylmalonic acidemia. Therefore, our study suggests that mitochondria dysfunctions caused by MMA in cells during the differentiation process may play an important role in the neuronal development delay which could be related to the neurologic symptoms as the cognitive deficit presented by some patients with methylmalonic acidemia [40]. Further studies would be necessary to determine the alterations on the functionality of neuronal cells after cellular bioenergetics impairments caused by MMA.

## 5. CONCLUSIONS

In this study, we provide evidence that MMA compromised cellular bioenergetics by reducing respiratory parameters in undifferentiated and RA-differentiated SH-SY5Y cells suggesting that mitochondrial dysfunction may play an important role in the pathogenesis of methylmalonic acidemia. Supporting this notion, neuronal differentiation was compromised by MMA exposition in the early and late stages of differentiation in SH-SY5Y cells which may be related to the mitochondrial dysfunction reported here. Altogether, this evidence contributes to understanding the neurological alterations found in the patients.

Our findings may contribute to the exploration of altered molecular markers in the presence of elevated concentrations of MMA as SYP and ENO2 genes may consist of a novel and interesting approach to monitor the evolution of neurologic alterations in patients within this disease.

## ABBREVIATIONS

AdoCbl: 5’-desoxyadenosylcobalamin
CCCP: carbonyl cyanide 3-chlorophenylhydrazone
cDNA: complementary DNA
CNS: central nervous system
DMEM: Dulbecco’s modified Eagle’s medium
DMSO: dimethyl sulfoxide
ENO2: neuron-specific enolase
FBS: fetal bovine serum
HPRT: hypoxanthine-guanine phosphoribosyltransferase
MMA: methylmalonic acid
MTT: methyl-thiazolyl diphenyl tetrazolium bromide
PBS: phosphate buffer saline
qPCR: quantitative PCR
RA: retinoic acid
ROX: residual oxygen consumption
SYP: synaptophysin.

